# Aberrant epithelialization: A plausible factor for the development of endometrial polyps

**DOI:** 10.1101/2025.08.18.670791

**Authors:** Amruta D. S. Pathare, Ankita Lawarde, Katrin Täär, Sergio Vela Moreno, Apostol Apostolov, Vijayachitra Modhukur, Darja Tarassova, Alberto Sola-Leyva, Andres Salumets, Merli Saare, Maire Peters

## Abstract

Endometrial polyps (EPs) are localized overgrowths of endometrial glands and stroma, common in reproductive-age and postmenopausal women, and can cause abnormal uterine bleeding and infertility. Here, we investigated the cellular heterogeneity and molecular mechanisms of EPs by integrating bulk and single-cell RNA sequencing (scRNA-seq) of EPs and adjacent endometrial tissues (adENs) from 12 women. Bulk RNA-seq revealed high transcriptional similarity, with few differentially expressed genes including upregulated KMT2B and DLEC1 and downregulated *COL9A1* and *RAB3C*. ScRNA-seq identified eight major cell clusters, such as stromal, epithelial, endothelial, immune, perivascular, macrophage, B, and ciliated cells. Pseudotime analysis showed aberrant stromal-to-epithelial transitions in EPs, marked by *MECOM* and *EYA2* intermediate clusters, indicating incomplete epithelial maturation. These altered differentiation trajectories may disrupt perivascular and endothelial cell development, contributing to abnormal vascular remodeling in EPs, despite minimal overall transcriptomic changes compared with adENs.

## Introduction

Endometrial polyps (EPs) are considered hyperplastic growths of endometrial stroma and glands ^1^. The prevalence of EPs ranges from 10% to 24%, and they are commonly observed in both reproductive-age and postmenopausal women ^1,2^. Although most EPs are asymptomatic, in some cases, they are accompanied by abnormal vaginal bleeding ^3^, dysmenorrhea and infertility ^4^, eventually affecting daily quality of life. EPs can also be associated with differential diagnosis and concomitant intrauterine pathologies like endometrial hyperplasia, adenomyosis, endometriosis, chronic endometritis, and, in rare cases, endometrial cancer with a low frequency of 0.8%-3.5% ^1,4–7^. In the presence of EPs, the endometrial tissue can display a spectrum ranging from normal cycling endometrium to simple or complex hyperplasia.

Development of EPs is reported to be hormone-dependent, primarily driven by estrogen, as indicated by increased immunohistochemical expression of estrogen and progesterone receptors in the glandular epithelium ^8,9^. Further, the increase in localised protein expression of BCL-2 in EPs during the proliferative phase suggests involvement of dysregulated cellular proliferation and apoptosis mechanisms ^10^. The activated mast cells and their oversecretion have been implicated in the pathogenesis of EPs, promoting enhanced angiogenesis and thickening of blood vessels in EPs ^11–13^. The significance of transcriptional regulator WT1 in the proliferation of mast cells has also been recently reported in the first single-cell RNA sequencing (scRNA-seq) study of EPs ^14^. EPs can affect fertility by altering the expression of endometrial receptivity-associated genes such as *HOXA10, HOXA11, and* prokineticin gene family ^2,15^. It can also cause mechanical hindering of embryo implantation based on the size and location in the fundus ^12,16^. Most of the reported mechanisms for the pathogenesis of EPs were based on targeted genes or protein expression, and thus, the precise mechanisms underlying EPs remain elusive.

Hysteroscopic polypectomy is currently the only treatment for EPs, highlighting the need to explore their molecular mechanisms to develop targeted drug therapies, especially for recurrent or symptomatic cases. Additionally, the transcriptome profile and cellular heterogeneity of EPs have been explored only in a few studies ^14,17^. In the current study, we used bulk and scRNA-seq to elucidate cellular heterogeneity and cell-specific transcriptomic alterations of EPs compared with adjacent endometrial tissue (adEN). Further, we explored the dynamic cellular transitions across the differentiation trajectory along with pseudotime analysis at a deeper resolution to uncover the factors contributing to the development of EPs. Our overall results suggest that EPs largely resemble normal endometrium but show subtle changes in genes linked to epithelial differentiation and tumorigenesis, suggesting protective mechanisms against malignancy. Altered epithelial differentiation may disrupt vascular remodelling in EPs. Further studies on epigenetics and functional models are needed to clarify these mechanisms.

## Results

The outline of the study is presented in Figure. 1. This study involved paired EPs and adEN samples from 12 women undergoing hysteroscopic polypectomy across the proliferative (n=9) and secretory (n=3) phases of the menstrual cycle (Figure 1a). Bulk RNA sequencing was performed on all samples (Figure 1b) including different comparison groups (Figure 1c and 1d), while scRNA-seq was performed on a subset of samples (five EPs and four adENs) (Figure 1e).

**Figure 1.**
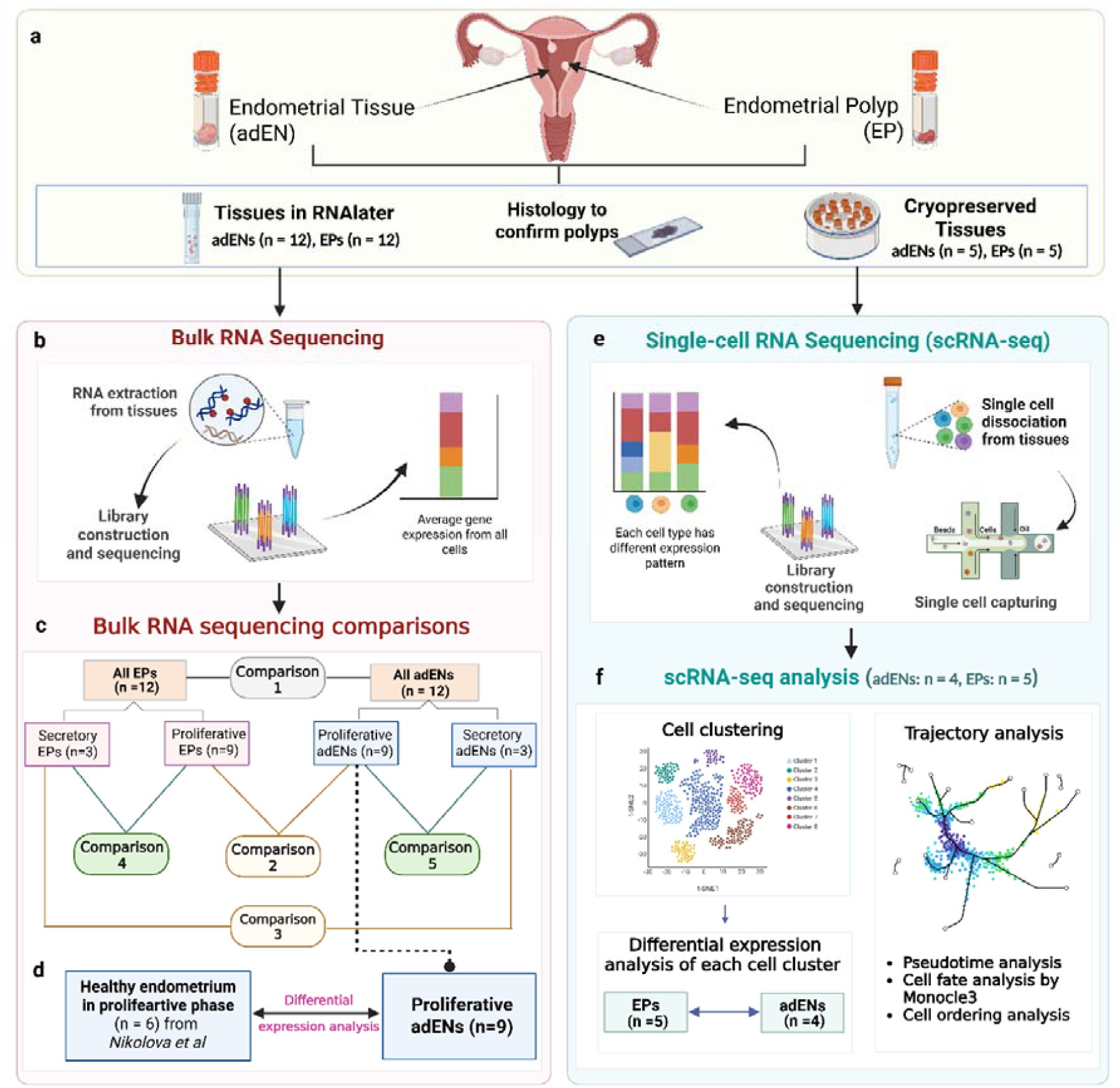
Outline of the study. **a.** Paired endometrial tissues (adENs) and polyps (EPs) were collected from 12 women in the proliferative phase (n = 9) and in the secretory phase (n = 3). Tissue samples were dissected and divided into three parts: used for histological evaluation, preserved in RNAlater stabilising solution for bulk RNA sequencing, and a subset of tissues (n = 5) were cryopreserved for single-cell RNA sequencing (scRNA-seq). **b.** Methodology of bulk RNA sequencing; RNA extraction, library preparation and sequencing were performed as per the manufacturer’s instructions. **c.** Transcriptome analysis from bulk RNA was performed between five groups; comparison 1: All EPs versus all adENs, comparison 2: EPs versus adENs in the proliferative phase, comparison 3: EPs versus adENs in the secretory phase, comparison 4: EPs in the secretory phase versus the proliferative phase, comparison 5: adENs in the secretory phase versus proliferative phase. **d.** adENs from proliferative phase were compared with gene expression dataset from healthy endometrium (n = 6) as published in Nikolova *et al*. **e.** A subset of cryopreserved tissues (adENs, n = 5 and EPs, n = 5) were subjected to single-cell dissociation, single-cell capturing by generation of gel beads-in-emulsion, library construction and sequencing as per the manufacturer’s instructions (one of the adEN sample was clogged during single-cell capturing and therefore excluded for further analysis). **f.** scRNA-seq data analysis was performed, including cell clustering, differential expression analysis and Pseudotime trajectory analysis. This figure is created using BioRender.com

### Whole-tissue transcriptome analysis of EPs and adENs

Bulk RNA sequencing analysis of EPs and adENs resulted in gene expression data of 16,580 genes after filtration. Based on the clustering analysis, the PCA plot of the overall gene expression did not reveal distinct groups of endometrial tissues and polyps, indicating that they are transcriptionally similar (Supplementary Figure 1). The differential expression analysis, irrespective of menstrual cycle phases (Figure 1c, comparison 1), revealed only two DEGs (Padj < 0.05), upregulated *KMT2B* (log_2_FC = 1.89, Padj = 3.08 x 10^-4^) and downregulated *COL9A1* (log_2_FC = −4.56, Padj = 9.03 x 10^-3^) in EPs, suggesting that EPs exhibit minimal transcriptional distinctions from the adENs. According to PCA analysis, two samples (P48 and H3) were identified as outliers (Supplementary Figure 1a). Although the overall clustering remained similar after removing the outliers (Supplementary Figure 1b), the differential expression analysis resulted in three additional downregulated genes in EPs; *RAB3C* (log_2_FC = −5.42, Padj = 0.02), *GPR22* (log_2_FC = −3.02, Padj = 0.02) and immunoglobulin kappa variable gene *IGKV1-33* (log_2_FC = −5.38, Padj = 0.03).

Furthermore, we performed a comparison of EPs versus adENs separately in the proliferative and secretory phases (Figure 1c, comparisons 2 and 3, respectively). In the proliferative phase (8 EPs vs 9 adENs), we identified the same three DEGs *KMT2B* (log_2_FC = 1.93, Padj = 0.022)*, COL9A1* (log_2_FC = −4.89, Padj = 0.022), and *RAB3C* (log_2_FC = −6.37, Padj = 0.008) in EPs exhibiting similar gene expression changes as in the previous comparison, and in addition, a significant upregulation of the *DLEC1* gene (log_2_FC = 3.98, Padj = 0.008). In the secretory phase, the comparison of EPs with adENs (n = 3) revealed three DEGs, *XPNPEP3* (log_2_FC = −2.14, Padj = 9 x 10^-6^)*, C9orf129* (log_2_FC = −6.2, Padj = 0.007), and *SLC25A21* (log_2_FC=-3.66, Padj = 0.028). Among them, C9orf129 is a pseudogene, whereas two protein-coding genes, *SLC25A21* and *XPNPEP3*, are involved in mitochondrial functions.

Next, we compared gene expression profiles of EPs and adENs across the proliferative and secretory phases by performing menstrual phase-specific comparisons: secretory EPs vs. proliferative EPs, and secretory adENs vs. proliferative adENs (Figure 1c, comparisons 4 and 5, respectively). The comparison of EPs in different phases revealed 88 DEGs (Supplementary Data 1), with the highest FC in the secretory EPs for protein-coding genes *PLA2G4F* (log_2_FC = 8.4, P = 0.001) and *KHDRBS2* (log_2_FC = −4.25, P = 0.012). The comparison of proliferative and secretory phase adENs showed 232 DEGs (Figure 1c, Supplementary Data 2), with maximum upregulation of *CYP26A1* (log_2_FC = 7.7, P = 0.004) and downregulation of *IGHV1-69* (log_2_FC = −9.3, P = 0.005) in the secretory phase adENs samples. As expected, secretory phase adENs showed overexpression of secretory phase-specific endometrial genes, including *MT2A, OFD1, LRRC1, KCNJ2, ADAMTS8, BIRC3, SLC15A2,* genes related to secretoglobin (*SCGB1D4* and *SCGB1D2*) and *MUC1* ^18,19^. Amongst these DEGs, upregulation of *SCGB1D4, SCGB1D2, and ADAMTS8*, whereas downregulation of *MMP7*, were overlapping in secretory EPs and adENs.

Enrichment analysis of secretory phase DEG sets in both EPs and adENs revealed the most relevant processes related to cellular response to stress and metal ion response with upregulated genes (*MT1G*, *MT1F* and *MT1X*), including metallothionein family members known for protecting against DNA damage and apoptosis from heavy metal ions ^20,21^ and play a vital role to achieve endometrial receptivity ^19^ (Supplementary Figure 2).

### Comparison of bulk transcriptomics of endometria from EP patients and healthy women

The comparison of adENs of women with EPs and healthy women in the proliferative phase revealed 18 DEGs (five upregulated and 13 downregulated), of which 12 were protein-coding genes (Table 1). The gene *CHST8*, involved in the biosynthesis of carbohydrates and hormones, was most upregulated (log_2_FC = 5.6) in endometrium from patients with EP, whereas peptidase inhibitor 3, *PI3* was the most downregulated (log_2_FC = −9.1) protein-coding gene.

**Table 1:**
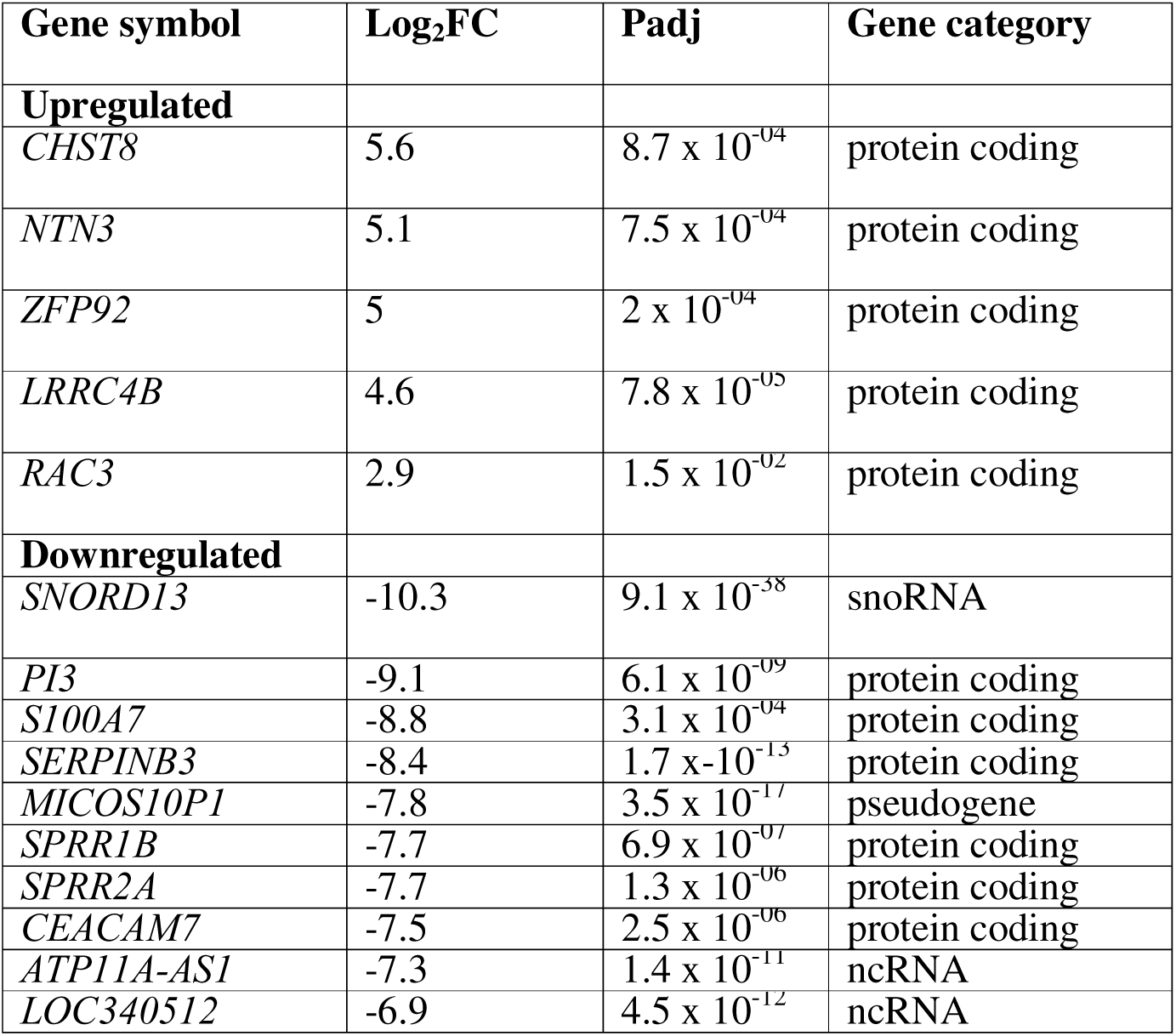
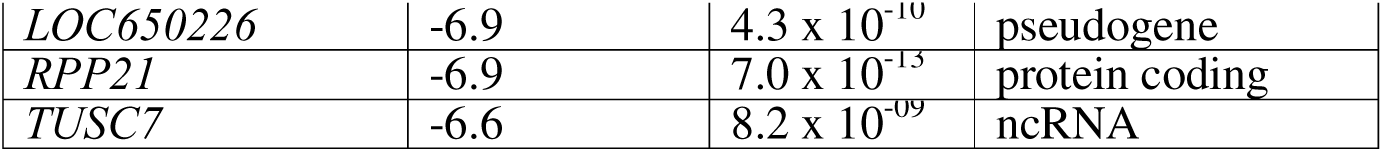
Dysregulated genes in the adjacent endometrium from women diagnosed with endometrial polyps compared to endometrium from healthy women. Abbreviations: FC: Fold change, Padj: adjusted P value.

### ScRNA-seq of EPs reveals a similar cellular composition to the adjacent endometrium

A total of 16,892 cells were obtained from scRNA-seq of four adENs and five EPs. One of the adEN samples was clogged during GEM generation and was excluded from further processing. After the quality control analysis and filtration of doublets, 7,011 cells from adENs and 8,075 cells from EPs were processed for further analysis. The details of scRNA-seq and quality control parameters are given in Supplementary Table 1.

The cells from the EPs and adENs were initially grouped into 21 clusters (Figure 2a), which were annotated based on the expression of cell-specific gene markers and regrouped into 10 major cell populations of stromal, epithelial, endothelial, immune, perivascular, macrophage, B cells, ciliated and two unknown clusters (Figure 2b and 2c). Unknown clusters include cells expressing genes not specific to any particular cell type. Of which, unknown cluster 2 shared few markers with endometrial cycling cells, such as *AGR2* and *KRT19* ^22^. In unknown cluster 1, which is an epithelial-like intermediate cluster, the majority of the cells were exhibited in EPs but absent in adENs (Figure 2c). Although differences in cell-type proportions were apparent between EPs and the adENs across clusters, these variations did not reach statistical significance (Figure 2d). The differential expression analysis between each cluster of EPs versus adENs showed 3 DEGs in the perivascular cluster: in EPs *BNC2* (log_2_FC = −3.7, Padj = 0.03) and *LINC01060* (log_2_FC = −4.6, Padj = 0.03) were downregulated, whereas *CSMD1* (log _2_FC = 3.3, Padj = 0.03) was upregulated (Figure 2e). The remaining clusters did not reveal a significant DEGs between the two groups, indicating that EPs possess a similar transcriptome to adENs.

**Figure 2.**
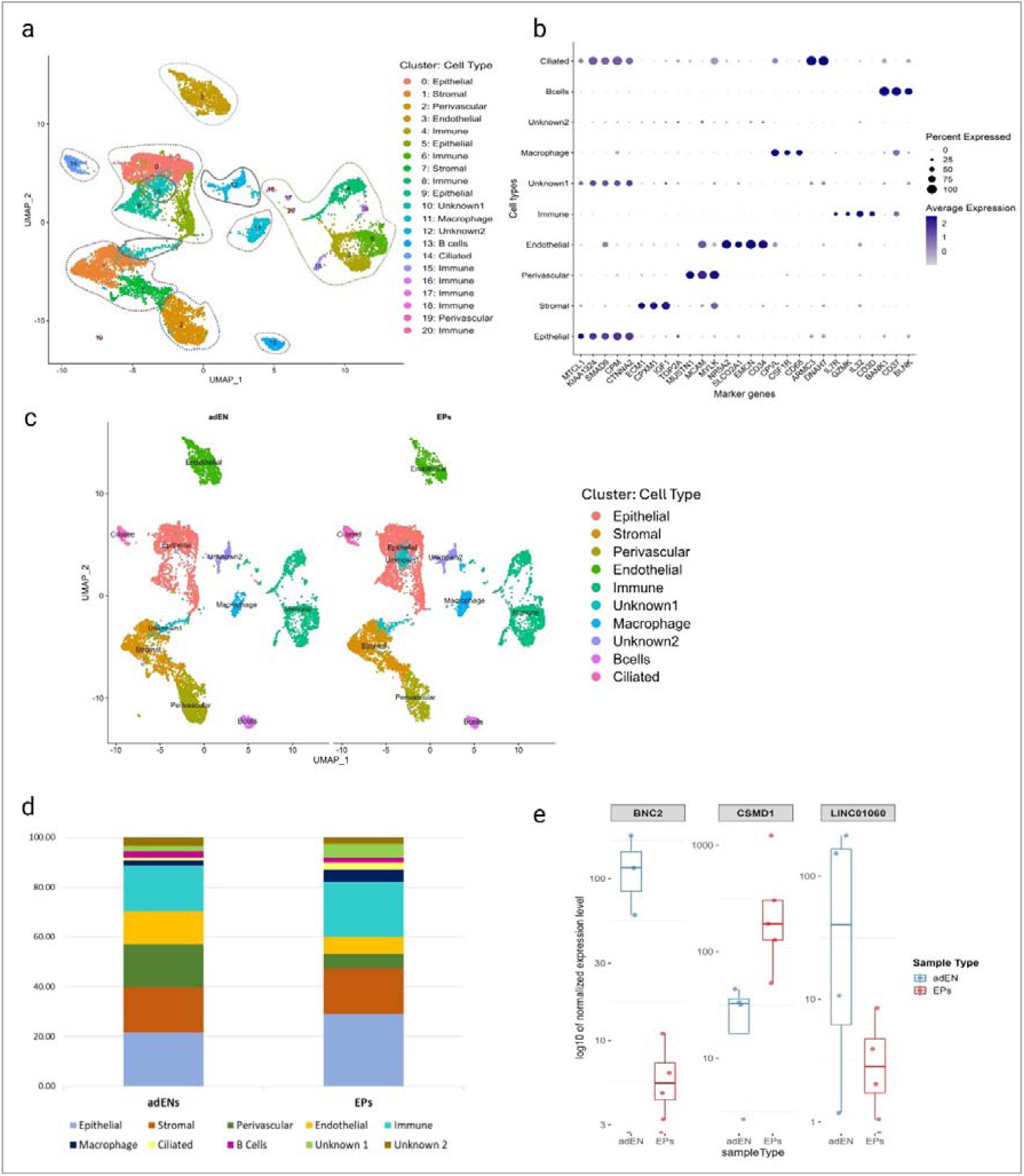
Cell clustering and differential gene expression of endometrial polyps (EPs) and adjacent endometrium (adENs) based on single cell RNA sequencing. **a.** Twenty-one different cell clusters were obtained in adENs and EPs at 0.4 resolution. The cell clusters were merged based on unique gene markers in each cluster. **b.** The unique cell markers used for cell cluster annotation of the 10 major populations. **c.** The merged cell clusters exhibited ten major cell populations which were subjected to further analysis. **d.** Difference in the cell percentage of each cluster in adENs and EPs did not show any statistical significance. **e.** Three genes *(BNC2, CSMD1, LINC01060)* were differentially expressed in the perivascular cluster of EPs compared to adENs.

### Trajectory and pseudotime analysis revealed aberrant and incomplete epithelialization in EPs

In EPs and adENs, the trajectory analysis revealed three paths arising from stromal cells and transitioning through unknown clusters 1 and/or 2 that led to different cell types, perivascular, endothelial, and ciliated cells (Figure 3a and 3b. Stromal cells have the lowest pseudotime, serving as the root of the trajectory in both EPs and adENs (Figure 3c and 3d. In both adENs and EPs, the first trajectory (path 1) represented the transition from stromal to perivascular cells. However, the sequence of cell fate transitions differed between conditions: in adENs, the stromal-to-endothelial and stromal-to-ciliated transitions appeared as path 2 and path 3, respectively, whereas in EPs, the stromal-to-ciliated transition occurred earlier (path 2), followed by the stromal-to-endothelial transition (path 3) (Figure 3a and 3b).

**Figure 3.**
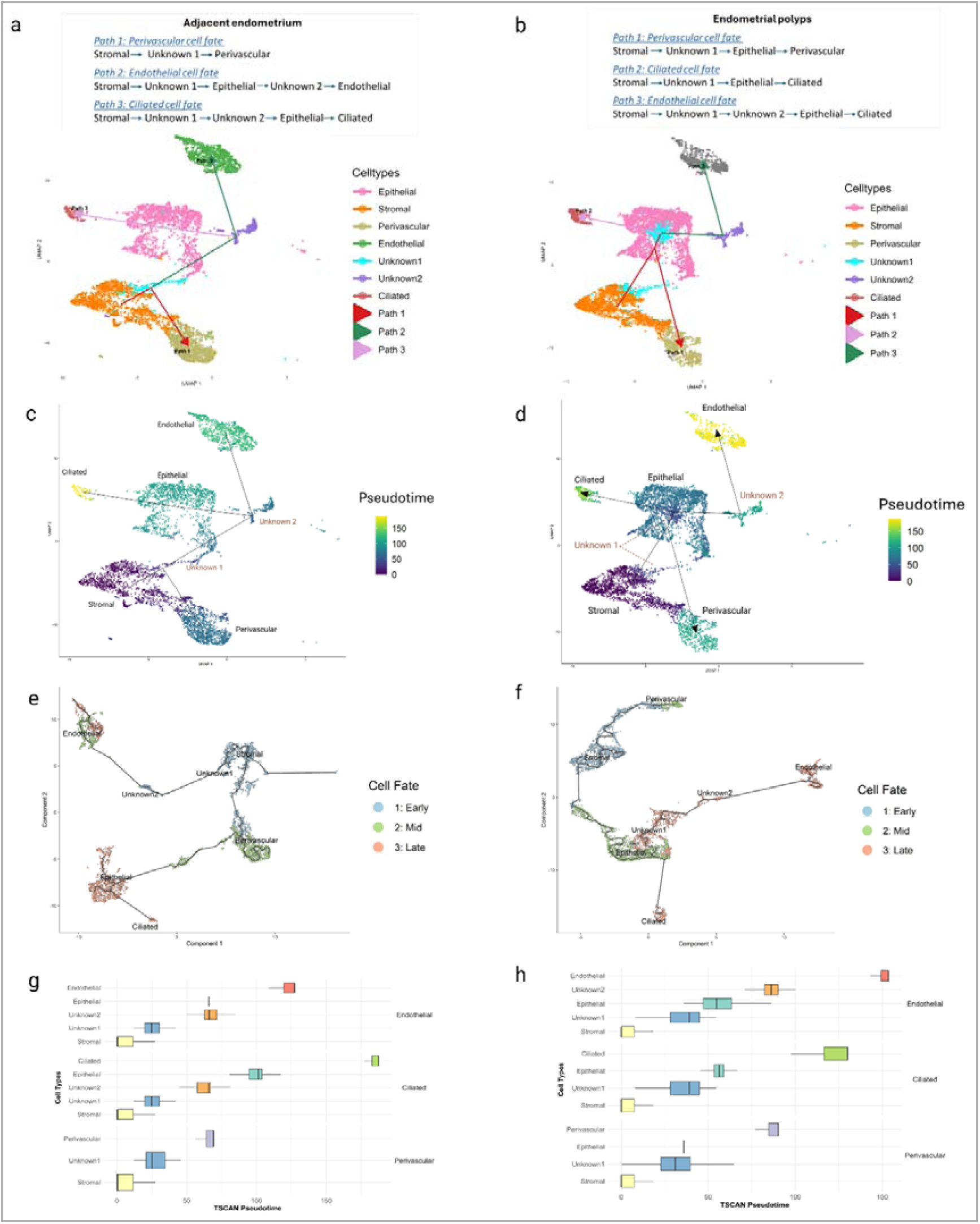
Trajectory and Pseudotime analysis of endometrial polyps (EPs) and adjacent endometrium (adENs). **a.** Reconstructed trajectories showed three different cell fates, such as perivascular, endothelial, and ciliated, in adENs based on single-cell RNA sequencing analysis, **b.** Pseudotime analysis of each of the three different cell fates in adENs, emphasising that stromal cells were the root of the trajectories, **c.** Cell fate analysis using Monocle3, validating the transcriptional transition states based on pseudotime in adENs, **d.** Cell ordering analysis across all three cell fates in adENs. **e.** Reconstructed trajectories showed three different cell fates, such as perivascular, endothelial, and ciliated, in EPs based on single-cell RNA sequencing analysis, **f.** Pseudotime analysis of each of the three different cell fates in EPs, indicating that stromal cells were the root of the trajectories, **g.** Cell fate analysis using Monocle3, validating the transcriptional transition states based on pseudotime in EPs, **h.** Cell ordering analysis across all three cell fates in EPs.

EPs exhibited an additional distinct trajectory, where stromal cells transitioned into epithelial cells via an unknown 1-cell cluster in all three cell fates. In contrast, in the adENs, this branching point is not observed, whereas the stromal to epithelial transition was observed only in the endothelial and ciliated paths. Moreover, the pseudotime of epithelial cells in EPs (Figure 3c) is markedly shorter compared to adENs (Figure 3d, showing a late transcriptional stage in adENs (Figure 3e), whereas a mid-stage in EPs (Figure 3f), suggesting impaired and incomplete epithelial maturation in EPs. This was further evidenced by cell ordering analysis based on pseudotime of ciliated cell fate (3g and 3h).

Furthermore, in mesenchymal-to-perivascular transition (MPT) in EPs, intermediate clusters, such as unknown 1 and epithelial clusters, exclusively involve cells with significantly high expression of *MECOM* (FDR = 9.9 x 10^-290^) and *EYA2* (FDR = 6.5 x 10^-290^) genes (Supplementary Data 3). These genes are subsequently repressed in the terminal perivascular cell cluster, indicating their specific enrichment in transitional states (Figure 4a). In contrast, in the adENs, *MECOM* (FDR = 6.2 × 10[²[) *and EYA2* (FDR = 0.0004) expression remains low throughout the MPT trajectory (Figure 4a, Supplementary Data 4), consistent with the absence of a substantial intermediate cluster toward the perivascular fate. Similarly, in the endothelial trajectory, specific expressions of epithelial markers such as *RHEX* (FDR = 2.54 x 10^-308^) and *CTNNA2* (FDR = 2.54 x 10^-308^) genes were enriched in transitional intermediate clusters in EPs, which were absent in adENs (Figure 4b, Supplementary Data 5 and 6). Based on Monocle3 trajectory analysis, all these genes showed significant Moran’s I test statistics, having FDR < 0.05 in both adENs and EPs, indicating consistent expression across the trajectory (Supplementary Data 7 and 8).

**Figure 4:**
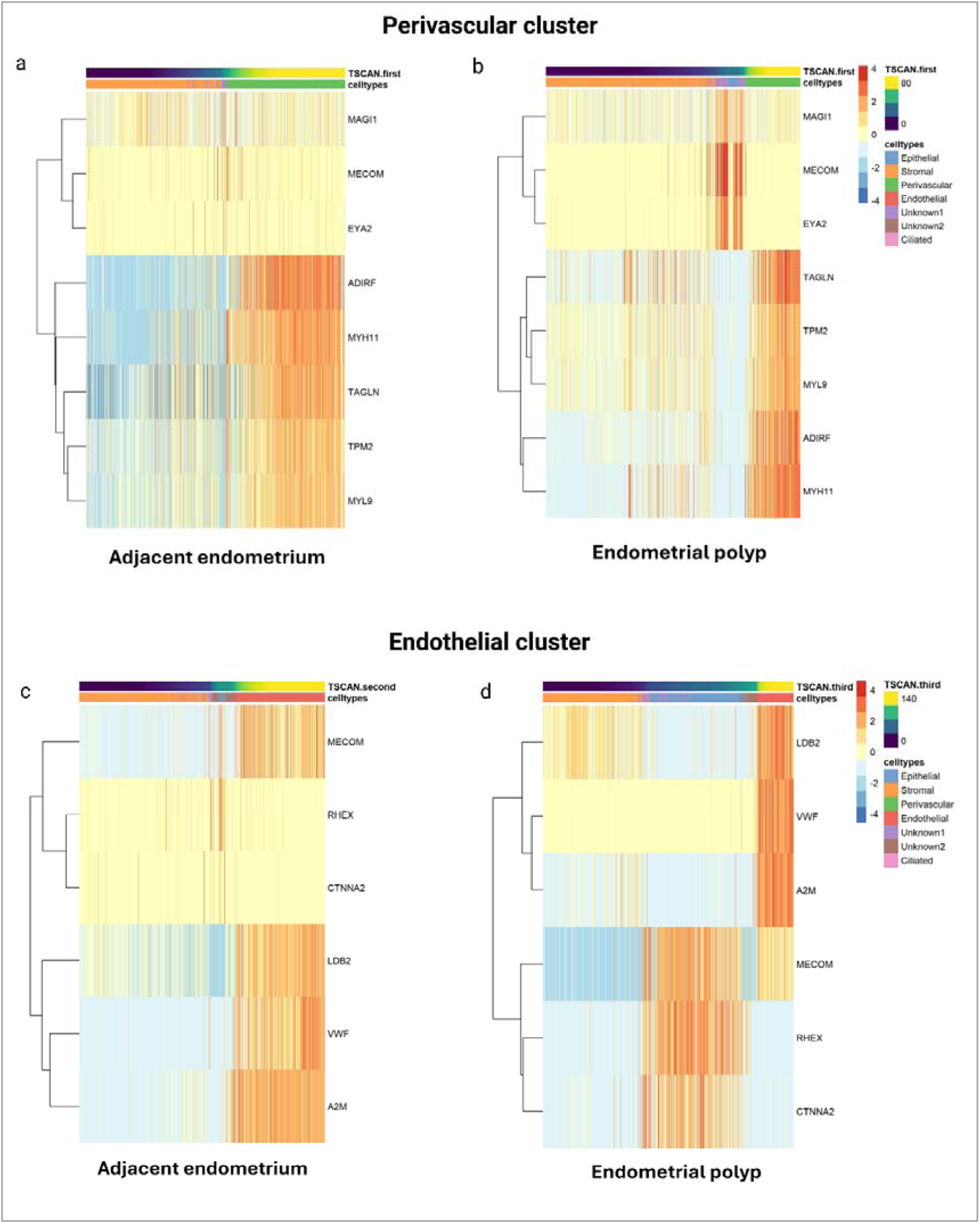
Heatmaps of pseudotime analysis across perivascular and endothelial clusters for selected genes having significant Moran 1 statistics in adjacent endometrium (adENs) and endometrial polyps (EPs). **a.** Differences in gene expression across perivascular trajectories in adENs (genes *MECOM, MAGI1* and *EYA2* showing altered expression in intermediate clusters and genes *ADIRF, MYH11, TAGLN TPM2* and *MYL9* being the signature for perivascular cells), **b.** Differences in expression of genes mentioned in Figure 4a across perivascular trajectories in EPs, **c.** Differences in gene expression in endothelial trajectories in adENs (genes *RHEX* and *CTNNA2* showing altered expression in intermediate clusters and genes *MECOM, LDB2, VWF* and *A2M* being the signature for endothelial cells), **s.** expression of genes mentioned in Figure 4c across endothelial trajectories in EPs.

## Discussion

EPs are a common endometrial pathology that affects the quality of women’s lives. Despite their high prevalence, the alterations in molecular mechanisms accountable for these endometrial outgrowths are not widely explored.

Our whole tissue bulk transcriptomic analysis showed that the gene expression profiles of EPs and adENs had only minimal differences. In the proliferative phase, the expression of *KMT2B*, *COL9A1, RAB3C* and *DLEC1* genes in EPs is highlighted, amongst which *COL9A1* and *RAB3C* were downregulated. *COL9A1*, involved in extracellular matrix (ECM) organisation, has been linked to endometrial receptivity and unexplained infertility, regulated via estrogen receptors ^23^ and upregulated in both primary and metastatic endometrial tumours^24^. The *RAB3C* gene encodes a small GTPase involved in intracellular vesicle trafficking — a process implicated in epithelial neoplasms and metastasis ^25^. Notably, *RAB3C* contains progesterone receptor binding sites and exhibits differential methylation in the extended promoter region of normal endometrial stromal fibroblast cells following treatment with estradiol and progesterone, indicating its regulatory role during decidualization ^26^. Although not directly linked to EPs, both genes are associated with endometrial abnormalities and tumour progression. While EPs possess the potential for malignant transformation, such progression is considered rare, with the overall prevalence of malignancy in EPs reported to be low ^27^ and most EPs remain benign. Therefore, our results showing downregulation of these genes in EPs may reflect a protective mechanism against tumorigenesis. The up-regulated *DLEC1*, a tumour suppressor gene, is involved in cell cycle regulation by inducing G1 phase arrest by inhibiting the NF-Kappa B inflammatory pathway ^28–30^. Similarly, the *KMT2B* gene has been associated with chromatin modelling and methylation in human cancers, including endometrial carcinoma ^31^. Given that both *DLEC1* and *KMT2B* were upregulated in EPs in our study, future research should explore the epigenetic mechanisms that may prevent them from developing into tumours.

Due to the transcriptomic similarity between EPs and adENs, we speculated that changes may have already occurred in the endometrium adjacent to the polyps. However, the comparison of adENs from our dataset and between published datasets of healthy endometrium ^32^ showed only 13 protein-coding DEGs. Amongst which, the *RPP21* gene, which is involved in tRNA processing and maturation and is ubiquitously expressed in all cell types of the endometrium ^22,33^. The *PI3* gene is expressed in neutrophils of human endometrium, specifically in the menstrual cycle phase ^34^. It is known for its anti-inflammatory and protease-inhibitory functions, which help protect the endometrial epithelial layer from inflammation, protease-mediated injury, and menstrual shedding-associated damage ^34,35^. Similarly, *SPRR2A* is involved in an antibacterial defence mechanism and protection against other environmental and extracellular damages. Its expression is estrogen-dependent and upregulated in the luminal epithelium of mouse endometrium during the oestrous cycle ^36^. However, its expression in the human endometrium has not been directly proven. In our dataset, downregulation of *RPP21*, *PI3 and SPRR2A* genes in adENs may indicate dysregulation in maintaining the normal functioning of the endometrium.

Furthermore, cellular heterogeneity by scRNA-seq revealed 10 major cell clusters shared between EPs and adENs, with no significant differences in their cell proportions; however, the analysis highlighted the involvement of perivascular cells in the EPs development with significant differences in expression of *BNC2, CSMD1* and *LINC01060* genes. The exact role of perivascular cells in EP pathogenesis is currently unclear. In the normal endometrium, perivascular cells, including pericytes, have a mesenchymal origin and are located near blood vessels. Endometrial perivascular cells play a role in regenerating capacities and repair, endometrial vascularisation and immunomodulation ^37–39^. Pericytes are also important in the formation of fibroblast-like cells and in maintaining the ECM ^38^. Any disturbances in the process of angiogenesis or degradation of the ECM would result in aberrant vasculature and abnormal bleeding ^38^. Thus, as EPs are characterised by the thickening of blood vessel walls, the DEGs in the perivascular cluster may offer insights into the mechanisms underlying endometrial tissue overgrowth, aberrant vascularisation, and the EPs formation. *BNC2* is a novel regulator of ECM composition and degradation, involved in epithelial-mesenchymal transition (EMT), and contributing to cell migration and invasion ^40,41^ whereas *CSMD1* is a tumour suppressor gene ^42^. Further, the *LINC01060* gene is again associated with EMT-related long non-coding (lnc)RNA and in our dataset, it was significantly downregulated in the perivascular cluster of EPs. It is involved in the ERK5 MAPK signalling pathway, which has been previously reported in the regulation of vascular endothelial growth factor (VEGF) expression for embryonic angiogenesis ^43,44^. Since *LINC01060* shares a common pathway for regulating angiogenesis, further studies in EPs, focusing on the lncRNA and even miRNA for regulating gene expression, would offer valuable insights. Furthermore, since the EMT-mesenchymal–epithelial transition (MET) processes are important physiological phenomenon that maintains endometrial functioning in terms of regeneration and re-epithelialization, their role in polyp development can be explored in detail ^45,46^. This was further supported by Lin *et al.*, who demonstrated a two-fold increase in the cross-sectional area of the glandular epithelium in the EP group compared to the eutopic endometrium ^14^.

The trajectory analysis of scRNA-seq further highlighted the importance of perivascular and epithelial cells. MPT trajectory of EPs exhibited an aberrant differentiation pattern compared to the adENs. In adENs, stromal cells (showing mesenchymal origin) directly differentiate into perivascular cells supporting normal vascular remodelling ^47^. In contrast, in EPs, the trajectory revealed an atypical path in which stromal cells transit through an epithelial cell cluster before differentiating into perivascular cells, indicating a disrupted MPT that may contribute to the altered vascular remodelling. Notably, in this abnormal lineage progression, genes like *MAGI1, EYA2* and *MECOM* were highly and exclusively expressed in the transitional clusters, including epithelial cells. *MECOM* is a transcription factor linked to stemness and tumour development ^48,49^ and is normally expressed in glandular and luminal epithelium contributing to epithelial regeneration ^22,50^. Its overexpression has been implicated in endometrial cancer and endometriosis ^49,51^. *MAGI1* is a scaffold protein important for maintaining epithelial polarity and angiogenesis ^52^. *EYA2* is a critical EMT regulator and is involved in cell differentiation, DNA repair mechanisms and inducing metastasis ^53^. However, in our scRNA-seq data, these genes were not differentially expressed across cell clusters, indicating that their apparent involvement in the MPT trajectory arises solely from the presence of epithelial cells within the transitional path, which is a deviation from normal lineage progression unique to EPs. Similar to the MPT trajectory, the endothelial trajectory in EPs contains a larger population of epithelial-like cells compared to adENs. Epithelial markers such as *RHEX* and *CTNNA2* were expressed exclusively in intermediate clusters along this trajectory. *RHEX* is expressed in endometrial epithelial cells by scRNA-seq ^50^ and associated with erythroid progenitor expansion ^54^, while *CTNNA2* is involved in cell–cell adhesion ^55^. Because normal stromal-to-endothelial differentiation does not include an epithelial intermediate, the presence of these markers likely reflects an aberrant differentiation path with inappropriate epithelial-like states, which may contribute to disrupted endothelial lineage commitment in EPs. Overall, the consistent expression of these genes along pseudotime, supported by significant Moran’s I statistics, suggests that abnormal epithelialization in EPs affects both mesenchymal and endothelial differentiation paths. This may underlie the cellular plasticity, dedifferentiation, and defective vascular remodelling observed in EPs. Therefore, the altered pattern of epithelial differentiation, which is not captured by conventional cluster-based transcriptomic analysis, may play a role in the development of polyps.

Among the three trajectories analysed in EPs, the pseudotime for the epithelial cluster was shorter than in adENs, suggesting incomplete epithelial differentiation and potentially aberrant mesenchymal-to-epithelial transition (MET). This is further supported by the considerably shorter pseudotime of the ciliated cell trajectory in EPs compared to adENs. As ciliated cells arise through epithelial differentiation, this finding reinforces the notion of disrupted or premature epithelialization in EPs. Such aberrant and excessive epithelial commitment may explain the increased epithelial cell proportion and reduced perivascular and endothelial cell populations in EPs, although these differences were not statistically significant. Notably, the presence of thickened blood vessel walls in EPs would typically suggest increased perivascular and endothelial cells, which contradicts our data. This discrepancy may be explained by excessive epithelialization, where cells appear to become stalled or retained in an epithelial-like intermediate state, diverting them from progressing toward proper vascular differentiation. Taken together, these observations suggest that altered epithelial differentiation may impair vascular remodelling and contribute to the unique pathology of EPs.

The bulk RNA sequencing study by Chiu et al. ^17^ compared EPs and the surrounding endometrium in infertile women and identified 322 DEGs, mainly related to WNT signalling and muscle contraction pathways, which may contribute to excessive proliferation and impaired vascular development. Although none of these DEGs were detected in our study, the DEGs found in both studies were associated with the same processes - excessive cellular proliferation and altered vascularisation. The recent scRNA-seq study by Lin *et al*. highlighted the role of activated mast cells as a mechanism of polyp development ^14^. In our study, we did not observe a separate cluster of mast cells, nor did the significant DEGs overlap, which could be due to the smaller number of cells targeted per sample in our study. Also, due to the inclusion of a heterogeneous population of EPs patients in the present study, clinical characteristics such as type, size and number of EPs, the presence of infertility and recurrence of polyps can influence the molecular mechanism and causal biological processes involved in EP development. Furthermore, our scRNA-seq results are restricted to the proliferative phase, as secretory phase samples were not included in this analysis. Another limitation of current single-cell studies in EPs is the absence of a control group consisting of healthy endometrial samples from non-EP women. As a result, the scRNA-seq analysis may have missed key cell-specific gene expression differences associated with EPs.

To summarise, our study reveals that the transcriptomic profile of EPs closely resembles that of normal endometrial tissue, except for a few genes, amongst which some were previously shown to be expressed in endometrial epithelial cells and involved in methylation and tumorigenesis. This suggests the potential protective mechanisms that prevent progression of EPs toward tumorigenesis. However, future studies focusing on epigenetic factors, such as methylation, are essential to further elucidate the mechanism involved. ScRNA-seq and trajectory analysis further revealed an important finding that aberrant and altered epithelial differentiation in EPs may disrupt the normal trajectory for perivascular and endothelial cells, resulting in abnormal vascular remodelling in EPs. Further functional studies, including *in vitro* epithelial organoid models in EPs, would help to understand the underlying mechanisms.

## Methods

### Materials and methods Ethical Approval

The study was approved by the Research Ethics Committee of the University of Tartu, Estonia (approval No 361T-19), and written informed consent was obtained from all participants.

### Patient selection and sample processing

Paired EP and adEN samples were obtained from 12 women (aged 39.2 ± 6.9 years, body mass index (BMI) 24.3 ± 4.1kg/m^2^, mean ± standard deviation) who had undergone hysteroscopy and polypectomy at the Tartu University Hospital (Tartu, Estonia). Samples were collected from different time points of the menstrual cycle: nine samples from the proliferative phase (cycle day 7-12) and three from the secretory phase (cycle day 17-21). None of the women had received hormonal treatments for at least three months before the time of hysteroscopy. Additionally, a subset of this cohort (n = 5)from the proliferative phase was subjected to scRNA-seq, (age range of 35-49 years and BMI 20.1-25.1 kg/m^2^). During hysteroscopic surgery, EPs were excised, and endometrial tissue biopsy was collected, placed in a hypothermic preservation medium, HypoThermosol® FRS Preservation Solution (Merck, Germany) and transported to the laboratory within 24 hours. Next, adENs and EPs were cut into small pieces and divided into three parts (Figure 1a); 1) immediately fixed in 4% formaldehyde for histological evaluation; 2) stored in RNAlater™ Stabilization Solution (Thermo Fisher Scientific, USA) for bulk RNA sequencing; and 3) added into cryopreservation medium containing 1× Dulbecco’s Modified Eagle’s Medium (DMEM, Gibco, USA), 30% fetal bovine serum (FBS, Gibco, USA) and 7.5% Dimethyl Sulfoxide Hybri-Max (Sigma-Aldrich, USA). Tissue samples preserved in the cryopreservation medium were placed into Nalgene Cryo 1°C ‘Mr Frosty’ Freezing Container (Thermo Fisher Scientific, USA) overnight in a −80°C freezer and stored in liquid nitrogen until batch-wise processing for scRNA-seq. The tissues fixed with 4% formaldehyde were embedded in paraffin, and sections of 5µm were cut and stained with Hematoxylin and Eosin (H&E) for routine histological evaluation. Upon histologically evaluating the tissue, the pathologist conclusively identified the presence of EPs based on their characteristic features (Supplementary Figure 3).

### Bulk RNA sequencing

To evaluate transcriptomic changes in the EPs compared to adENs, we performed bulk RNA sequencing on a total of 24 samples collected from 12 women (Figure 1b). Total RNA was extracted from the tissues using the miRNeasy Tissue advanced mini kit (Qiagen, Germany) according to the manufacturer’s instructions. Bulk RNA libraries were performed using 500 ng total RNA by TruSeq Stranded mRNA library kit and TruSeq RNA CD Index Plate according to the manufacturer’s (Illumina, USA) instructions. The libraries were sequenced on a NextSeq 1000 (Illumina, USA) sequencer using single-end 80 bp setting as per the manufacturer’s instructions, resulting in an average of 15 million reads per sample. Differential gene expression analysis by DESeq2 method was performed using Illumina BaseSpace. Briefly, fastq files from the sequencing run were generated using BCL Convert app (v2.1.0), which was further processed by DRAGEN FASTQ Toolkit app (v1.0.0) to trim the adapter sequences and DRAGEN FastQC + MultiQC (v3.9.5) was used for quality control analysis. Further steps, such as alignment with hg38 Alt-Masked v2, normalization and differential expression analysis by DESeq2 were performed by DRAGEN RNA (v3.10.4). The differentially expressed genes (DEGs) between EP and adENs having log2 fold change (log_2_FC) > 1.0 and FDR P value < 0.05 were considered significant. Different comparison groups were analysed as shown in Figure 1c. Enrichment analysis for DEGs was performed using g:Prolifer web based tool ^56^.

### Integration of bulk RNA sequencing data of healthy endometrium

We performed differential expression analysis of adENs collected from EP patients (n=9) from our dataset with a gene expression dataset ^32^ from healthy endometrium (n=6) in the proliferative phase using R v4.4.1 (Figure 1d). We applied a pre-filtering step and retained genes with at least 10 raw counts in a minimum of 3 samples. The batch effect adjustment was performed by ComBat-Seq method which was implemented from R package (Bioconductor sva v3.36.0) ^57^.

### Tissue dissociation for scRNA-seq

To determine the cellular heterogeneity of EPs and corresponding transcriptome profile for each cell type, we performed scRNA-seq using 10x Genomics technology on EPs and adENs from five women in the proliferative phase. (Figure 1e). Cryopreserved tissues were thawed in a 37°C water bath and washed twice with a pre-warmed DMEM medium (Phenol red free, ccFBS 10% with penicillin-streptomycin, Corning USA) to eliminate cryoprotectant and excess blood. Enzymatic dissociation using 2.5mg/ml collagenase from Clostridium histolyticum (Sigma-Aldrich, USA); 0.25mg/ml DNase (AppliChem GmbH, Germany) and 10mg/ml Dispase II (Thermo Fisher Scientific, USA) in 10ml of the DMEM medium was performed for approximately 30 mins at 37°C on a rotating shaker. Undigested tissue fragments were settled by gravity and re-digested enzymatically for an additional 15 mins. The resulting single cell suspension was filtered (30-μm cell strainer), pelleted by centrifugation at 300×g for 5 mins, and treated with ammonium–chloride–potassium (ACK) lysis buffer (Gibco, USA), to remove red blood cells and following the manufacturer’s instructions. Live cells having viability of >90% were enriched, washed with 0.04% BSA (Sigma-Aldrich, USA) in Dulbecco’s (D)PBS (1×, Gibco, USA) on ice to eliminate ambient RNAs, and resuspended in 0.04% BSA in DPBS to attain a concentration of 700-1200 cells/μl as per the 10x Genomics protocol.

### Chromium 10x single cell capturing, library generation and sequencing

Gel Beads-in-emulsion (GEM) of single cell suspension was generated by 10x Chromium Controller. To target 5,000 cells per sample, the single cell suspension, single cell 3’ gel beads, and master mix for reverse transcription reagents were loaded onto a 10x Chromium microfluidic chip. The preparation of single cell suspension and library construction were performed using 10x Chromium Next GEM Single Cell 3′ reagent v3.1 kit (Dual index, 10x Genomics CG000315 Rev C) comprised of single cell 3’ gel bead kit, library construction kit and Chip G kit as per the manufacturer’s instructions (Figure 1e). Subsequent steps encompassed cell lysis, first-strand cDNA synthesis, amplification, and purification to yield barcoded full-length cDNA, which were conducted according to the manufacturer’s protocol. To mitigate batch effects, library preparation was performed simultaneously for all samples, ensuring consistency. Quality assessment of cDNA and single-cell libraries was performed using an Agilent 4150 Tape Station. The dual-indexed single-cell libraries were combined and subjected to paired-end sequencing on the NovaSeq PE150 platform, aiming for 35,000 reads per cell.

### Quality control, normalization, and cell clustering of scRNA-seq data

The scRNA-seq data from 10x Genomics was mapped to the reference genome, refdata-gex-GRCh38-2020-A (https://www.10xgenomics.com/). Count matrices were generated for individual samples using Cell Ranger software (v7.0.0).

The aligned scRNA-seq data was analysed using the CRAN package Seurat (v.5.3.0). During the data processing steps, cells were selected based on specific criteria: those with a minimum Unique Molecular Identifier (UMI) count exceeding 500 and cells expressing over 200 genes were retained. Cell types showing high complexity (greater than 0.80) were preserved, while cells with more than 15% mitochondrial reads were excluded from the analysis. Genes with zero counts across the dataset were eliminated, allowing only genes expressed in at least three cells to remain. Subsequently, each sample underwent normalization, and the CellCycleScoring function within the Seurat package was utilized to assign a cell cycle score. This score was based on the expression of markers associated with the G2/M and S phases, obtained from ^58^, as outlined in the Seurat package manual. The SCTransform function from Seurat package was employed to normalize the data ^59^, estimate the variance of raw filtered data, and identify the most variable genes. This process aimed to eliminate the effect of variations caused by mitochondrial expression and cell cycle phases. Subsequently, Principal Component Analysis (PCA) analysis was performed using SCTransformed samples, where the minimum number of PCs (n = 40) were identified. Using the resultant PC, we conducted Uniform Manifold Approximation and Projection (UMAP) clustering, as well as executed the FindNeighbors and FindClusters functions in Seurat with 0.1 resolution to delineate distinct clusters within the dataset. Further, the pre-processed data was subjected to the analysis using the DoubletFinder R package with default settings ^60^. Identified doublets were eliminated from each sample before the integration of all the samples. For integration purposes, we used 5,000 highly variable features common to all samples to identify shared subpopulations across the adENs and EPs. Canonical correlation analysis (CCA) was conducted to reveal common sources of variation between these two groups. In brief, CCA identifies significant sources of variation within the data, specifically those shared or conserved across various conditions or groups, utilising the 5,000 variable features. In the subsequent integration step, mutual nearest neighbours (MNN) were determined, and incorrect anchors (cells) were removed to integrate the samples across different conditions.

Following integration, the data was visualised through dimensionality reduction techniques such as PCA and UMAP. The FindClusters function in Seurat, with resolutions spanning from 0.2 to 1.4, was employed to identify distinct clusters within the integrated data. Finally, this integrated dataset with identified cell clusters visualised by the UMAP method with 0.4 resolution was used for further downstream analysis.

### Cell cluster annotations

To identify and annotate major cell types in the integrated dataset we used two distinct approaches. Initially, the FindAllMarkers function within the Seurat package was run by setting default parameters to detect DEGs associated with each specific identity class. Further, we compiled cell markers sourced from various references from the literature ^22,61^, cell marker database ^62^, and the scRNA-seq database obtained from the Human Protein Atlas to validate the identified cell markers ^50^. To shortlist the selection of marker genes, we applied the following criteria: log2FC >1, adjusted p-value (Padj) < 0.05, PCT.1 ≥ 0.7 (indicating higher expression of a marker in a specific cluster), and PCT.2 ≤ 0.3 (indicating lower expression of the same marker in other clusters) (Figure 1f). Statistical analysis was performed using chi-square and Fisher’s exact test to determine the cell proportions and the comparison of total cell numbers between adENs and EPs.

### Pseudotime and trajectory analysis

To investigate dynamic transcriptional changes underlying cellular progression among resident endometrial cell types, we performed trajectory analysis on all major clusters, excluding the immune cell cluster, which originates from a hematopoietic lineage and is transcriptionally distinct from the tissue-resident cell populations (Figure 1f).

The analysis was performed using the TSCAN R package (v1.42.0) ^63^, which implements a graph-based method based on minimum spanning trees (MSTs). All analyses were conducted using default parameters unless otherwise specified. The integrated Seurat object generated from prior steps was used as the input and was converted to a SingleCellExperiment (SCE) object to ensure compatibility with TSCAN functions. Separate trajectory analyses for EPs and adENs samples were conducted.

Trajectory inference was initiated by aggregating gene expression profiles across cells belonging to each cell-type-annotated cluster using the *aggregateAcrossCells* function from the scater package (v1.32.1). Cluster centroids were computed in the PCA space and subsequently used to construct a global trajectory via the *createClusterMST* function. The resulting MST illustrated the shortest Euclidean-distance connections among clusters within the reduced dimensional space.

Pseudotime values were assigned by projecting individual cells onto the MST structure. Cells were mapped to the nearest MST edges using the *mapCellsToEdges* function, and pseudotemporal ordering was derived using *orderCells* function. This enabled continuous reconstruction of cell-state transitions along inferred developmental trajectories, independent of prior clustering labels. In instances where cells were associated with multiple trajectory branches, pseudotime values were averaged to accommodate potential branching events.

To investigate transcriptional dynamics along the inferred trajectories, we identified genes whose expression patterns significantly varied with pseudotime. This was achieved using the *testPseudotime* function in TSCAN, which fits a natural spline to model gene expression as a non-linear function of pseudotime. An analysis of variance (ANOVA) was then applied to test whether the spline coefficients significantly deviated from zero, indicating dynamic regulation. Genes with low p-values (FDR < 0.05) were considered as key contributors to trajectory progression.

Trajectory inference was performed using the Monocle3 R package (v1.2.9) to complement the TSCAN-based analysis and to capture additional transcriptional dynamics. The integrated Seurat object generated from previous steps was used as the input. The object was converted into a cell_data_set (CDS) object using the *as.cell_data_set* function to ensure compatibility with the Monocle3 framework. Similar to TSCAN, separate trajectory analysis was conducted for EPs and adENs.

Following conversion to CDS object, the precomputed UMAP coordinates and cluster annotations from Seurat were retained for trajectory reconstruction. The trajectory graph was learned using Monocle3’s *learn_graph* function, which constructs a principal graph capturing cell state transitions within the low-dimensional embedding space.

To identify genes with dynamic expression patterns along the inferred trajectories, we applied the *graph_test* function, which uses Moran’s I statistic to assess spatial autocorrelation between gene expression and the trajectory graph structure. Genes with high spatial autocorrelation (q-value < 0.05, Benjamini–Hochberg corrected) were considered significantly associated with the trajectory and indicative of pseudotime-dependent transcriptional regulation. Cell fate states were assigned by dividing pseudotime values into quantile-based intervals corresponding to early, mid, and late stages, and each cell was categorized accordingly based on its pseudotime value. The differentially regulated genes were visualised using a heatmap, and all trajectories were visualised on UMAP projections. Cell cluster ordering obtained from TSCAN was visualized using a boxplot.

## Data availability

The raw data for bulk transcriptomics and single-cell RNA sequencing are available online under the GEO accession numbers GSE305280 and GSE305281, respectively.

## Authors’ roles

Authors A.D.S.P. and A. L. conducted the experiments, performed bioinformatic data analysis and interpretation, and drafted and revised the manuscript. K.T., M. S and M. P. supported the study with sample collection and coordination. V. M. and S. V. M. contributed to data analysis. A. A., D. T. and A. S. L. participated in experiments. A. D. S. P., A. L., A. S., M. S., and M. P. contributed to the conception of the study, critically reviewed and provided feedback on each version of the manuscript. All the authors approved the final version of the manuscript.

## Acknowledgements

We acknowledge the use of BioRender.com to create Figure 1. We would like to express our gratitude to all the women in Estonia who agreed to participate in the study.

## References

1. Tanos, V. et al. The management of polyps in female reproductive organs. Int J Surg 43, 7–16 (2017).

2. Özbey, G., Tuncay, G., Düz, S. A., Çiğremiş, Y. & Karaer, A. The Effect of Endometrial Polyp and Myoma Uteri on Fertility-Related Genes in the Endometrium. Reprod Sci 32, 728–737 (2025).

3. Vieira, M. D. C. et al. Endometrial Polyps: Update Overview on Etiology, Diagnosis, Natural History and Treatment. CEOG 49, 232 (2022).

4. Raz, N., Feinmesser, L., Moore, O. & Haimovich, S. Endometrial polyps: diagnosis and treatment options - a review of literature. Minim Invasive Ther Allied Technol 30, 278– 287 (2021).

5. Dias Da Silva, I., Wuidar, V., Zielonka, M. & Pequeux, C. Unraveling the Dynamics of Estrogen and Progesterone Signaling in the Endometrium: An Overview. Cells 13, 1236 (2024).

6. Levy, R. A., Kumarapeli, A. R., Spencer, H. J. & Quick, C. M. Cervical polyps: Is histologic evaluation necessary? Pathol Res Pract 212, 800–803 (2016).

7. Annan, J. J., Aquilina, J. & Ball, E. The management of endometrial polyps in the 21st century. The Obstetrician & Gynaecologist 14, 33–38 (2012).

8. Inceboz, U. S. et al. Hormone receptor expressions and proliferation markers in postmenopausal endometrial polyps. Gynecol Obstet Invest 61, 24–28 (2006).

9. Lopes, R. G. C. et al. Analysis of estrogen-and progesterone-receptor expression in endometrial polyps. J Minim Invasive Gynecol 14, 300–303 (2007).

10. Taylor, L. J., Jackson, T. L., Reid, J. G. & Duffy, S. R. G. The differential expression of oestrogen receptors, progesterone receptors, Bcl-2 and Ki67 in endometrial polyps. BJOG 110, 794–798 (2003).

11. Al-Jefout, M. et al. Novel finding of high density of activated mast cells in endometrial polyps. Fertil Steril 92, 1104–1106 (2009).

12. Nijkang, N. P., Anderson, L., Markham, R. & Manconi, F. Endometrial polyps: Pathogenesis, sequelae and treatment. SAGE Open Med 7, 2050312119848247 (2019).

13. Norrby, K. Mast cells and angiogenesis. APMIS 110, 355–371 (2002).

14. Lin, Y. et al. Single-cell sequencing reveals a regulatory role of WT1 in mast cell proliferation in endometrial polyps. The FASEB Journal 39, e70512 (2025).

15. Rackow, B. W., Jorgensen, E. & Taylor, H. S. ENDOMETRIAL POLYPS AFFECT UTERINE RECEPTIVITY. Fertil Steril 95, 2690–2692 (2011).

16. Yanaihara, A., Yorimitsu, T., Motoyama, H., Iwasaki, S. & Kawamura, T. Location of endometrial polyp and pregnancy rate in infertility patients. Fertil Steril 90, 180–182 (2008).

17. Chiu, C. S.-C., Yeh, L.-Y., Pan, S.-H. & Li, S.-H. Transcriptomic Analysis Reveals Intrinsic Abnormalities in Endometrial Polyps. International Journal of Molecular Sciences 25, 2557 (2024).

18. Díaz-Gimeno, P. et al. A genomic diagnostic tool for human endometrial receptivity based on the transcriptomic signature. Fertil Steril 95, 50–60, 60.e1–15 (2011).

19. Ruiz-Alonso, M., Blesa, D. & Simón, C. The genomics of the human endometrium. Biochim Biophys Acta 1822, 1931–1942 (2012).

20. Chiaverini, N. & De Ley, M. Protective effect of metallothionein on oxidative stress-induced DNA damage. Free Radic Res 44, 605–613 (2010).

21. Qu, W., Pi, J. & Waalkes, M. P. Metallothionein blocks oxidative DNA damage in vitro. Arch Toxicol 87, 311–321 (2013).

22. Marečková, M. et al. An integrated single-cell reference atlas of the human endometrium. Nat Genet 56, 1925–1937 (2024).

23. Klonos, E. et al. Endometrial changes in estrogen and progesterone receptor expression during implantation in an oocyte donation program. Exp Ther Med 20, 178 (2020).

24. Yadav, V. K. et al. Computational analysis for identification of the extracellular matrix molecules involved in endometrial cancer progression. PLoS One 15, e0231594 (2020).

25. Goldenring, J. R. A central role for vesicle trafficking in epithelial neoplasia: Intracellular highways to carcinogenesis. Nat Rev Cancer 13, 813–820 (2013).

26. Houshdaran, S. et al. Steroid hormones regulate genome-wide epigenetic programming and gene transcription in human endometrial cells with marked aberrancies in endometriosis. PLoS Genet 16, e1008601 (2020).

27. Adomaitienė, L. et al. Proliferation in Postmenopausal Endometrial Polyps—A Potential for Malignant Transformation. Medicina 55, 543 (2019).

28. Seven, D. et al. DLEC1 is not silenced solely by promoter methylation in head and neck squamous cell carcinoma. Gene 563, 83–86 (2015).

29. Kwong, J. et al. Candidate Tumor-Suppressor Gene DLEC1 Is Frequently Downregulated by Promoter Hypermethylation and Histone Hypoacetylation in Human Epithelial Ovarian Cancer. Neoplasia (New York, N.Y.) 8, 268 (2006).

30. Ying, J. et al. DLEC1 is a functional 3p22.3 tumour suppressor silenced by promoter CpG methylation in colon and gastric cancers. Br J Cancer 100, 663–669 (2009).

31. Momeni-Boroujeni, A. et al. Landscape of chromatin remodeling gene alterations in endometrial carcinoma. Gynecologic Oncology 172, 54–64 (2023).

32. Nikolova, M. et al. Coupling miR/isomiR and mRNA Expression Signatures Unveils New Molecular Layers of Endometrial Receptivity. Life 11, 1391 (2021).

33. GTEx Consortium. The GTEx Consortium atlas of genetic regulatory effects across human tissues. Science 369, 1318–1330 (2020).

34. Lee, S., Yoo, I., Cheon, Y. & Ka, H. Spatiotemporal expression and regulation of peptidase inhibitor 3 and secretory leukocyte protease inhibitor at the maternal-fetal interface in pigs. Anim Biosci 36, 1034–1043 (2023).

35. King, A. E., Critchley, H. O. D. & Kelly, R. W. Innate immune defences in the human endometrium. Reprod Biol Endocrinol 1, 116 (2003).

36. Hong, S. H. et al. Estrogen regulates the expression of the small proline-rich 2 gene family in the mouse uterus. Mol Cells 17, 477–484 (2004).

37. Tang, Y. et al. Human Endometrial Pericytes: A Comprehensive Overview of Their Physiological Functions and Implications in Uterine Disorders. Cells 13, 1510 (2024).

38. Li, S. & Ding, L. Endometrial Perivascular Progenitor Cells and Uterus Regeneration. J Pers Med 11, 477 (2021).

39. Gharanei, S. et al. Vascular Adhesion Protein-1 Determines the Cellular Properties of Endometrial Pericytes. Front Cell Dev Biol 8, 621016 (2020).

40. Bobowski-Gerard, M. et al. Functional genomics uncovers the transcription factor BNC2 as required for myofibroblastic activation in fibrosis. Nat Commun 13, 5324 (2022).

41. Orang, A. et al. Basonuclin-2 regulates extracellular matrix production and degradation. Life Sci Alliance 6, e202301984 (2023).

42. Wang, X. et al. CSMD1 suppresses cancer progression by inhibiting proliferation, epithelial-mesenchymal transition, chemotherapy-resistance and inducing immunosuppression in esophageal squamous cell carcinoma. Experimental Cell Research 417, 113220 (2022).

43. Li, L. et al. Transcriptome sequencing of endometrium revealed alterations in mRNAs and lncRNAs after ovarian stimulation. J Assist Reprod Genet 37, 21–32 (2020).

44. Sohn, S. J., Sarvis, B. K., Cado, D. & Winoto, A. ERK5 MAPK regulates embryonic angiogenesis and acts as a hypoxia-sensitive repressor of vascular endothelial growth factor expression. J Biol Chem 277, 43344–43351 (2002).

45. Owusu-Akyaw, A., Krishnamoorthy, K., Goldsmith, L. T. & Morelli, S. S. The role of mesenchymal-epithelial transition in endometrial function. Hum Reprod Update 25, 114– 133 (2019).

46. Kirkwood, P. M. et al. Single-cell RNA sequencing and lineage tracing confirm mesenchyme to epithelial transformation (MET) contributes to repair of the endometrium at menstruation. eLife 11, e77663 (2022).

47. Patterson, A. L. et al. Putative human myometrial and fibroid stem-like cells have mesenchymal stem cell and endometrial stromal cell properties. Hum Reprod 35, 44–57 (2020).

48. Masamoto, Y. [The role and regulation of EVI1 in normal hematopoiesis and hematopoietic malignancies]. Rinsho Ketsueki 65, 954–960 (2024).

49. Lou, M. et al. MECOM and the PRDM gene family in uterine endometrial cancer: bioinformatics and experimental insights into pathogenesis and therapeutic potentials. Molecular Medicine 30, 190 (2024).

50. Karlsson, M. et al. A single–cell type transcriptomics map of human tissues. Science Advances 7, eabh2169 (2021).

51. Sarsenova, M. et al. Endometriotic lesions exhibit distinct metabolic signature compared to paired eutopic endometrium at the single-cell level. Commun Biol 7, 1026 (2024).

52. Wörthmüller, J. & Rüegg, C. MAGI1, a Scaffold Protein with Tumor Suppressive and Vascular Functions. Cells 10, 1494 (2021).

53. Farabaugh, S. M., Micalizzi, D. S., Jedlicka, P., Zhao, R. & Ford, H. L. Eya2 is required to mediate the pro-metastatic functions of Six1 via the induction of TGF-β signaling, epithelial-mesenchymal transition, and cancer stem cell properties. Oncogene 31, 552– 562 (2012).

54. Verma, R. et al. RHEX, a novel regulator of human erythroid progenitor cell expansion and erythroblast development. J Exp Med 211, 1715–1722 (2014).

55. Fanjul-Fernández, M. et al. Cell–cell adhesion genes CTNNA2 and CTNNA3 are tumour suppressors frequently mutated in laryngeal carcinomas. Nat Commun 4, 2531 (2013).

56. Raudvere, U. et al. g:Profiler: a web server for functional enrichment analysis and conversions of gene lists (2019 update). Nucleic Acids Research 47, W191–W198 (2019).

57. Zhang, Y., Parmigiani, G. & Johnson, W. E. ComBat-seq: batch effect adjustment for RNA-seq count data. NAR Genomics and Bioinformatics 2, lqaa078 (2020).

58. Tirosh, I. et al. Dissecting the multicellular ecosystem of metastatic melanoma by single-cell RNA-seq. Science 352, 189–196 (2016).

59. Hafemeister, C. & Satija, R. Normalization and variance stabilization of single-cell RNA-seq data using regularized negative binomial regression. Genome Biology 20, 296 (2019).

60. McGinnis, C. S., Murrow, L. M. & Gartner, Z. J. DoubletFinder: Doublet Detection in Single-Cell RNA Sequencing Data Using Artificial Nearest Neighbors. Cell Systems 8, 329–337.e4 (2019).

61. Garcia-Alonso, L. et al. Mapping the temporal and spatial dynamics of the human endometrium in vivo and in vitro. Nat Genet 53, 1698–1711 (2021).

62. Hu, C. et al. CellMarker 2.0: an updated database of manually curated cell markers in human/mouse and web tools based on scRNA-seq data. Nucleic Acids Res 51, D870– D876 (2023).

63. Ji, Z. & Ji, H. TSCAN: Pseudo-time reconstruction and evaluation in single-cell RNA-seq analysis. Nucleic Acids Res 44, e117 (2016).

