## Supplementary Figure 1 for "Aberrant epithelialization: A plausible factor for the development of endometrial polyps"

**Supplementary Figure 1: Principal component analysis of endometrial polyps (EPs) and adjacent endometrium (adENs) irrespective of the menstrual cycle phases.**

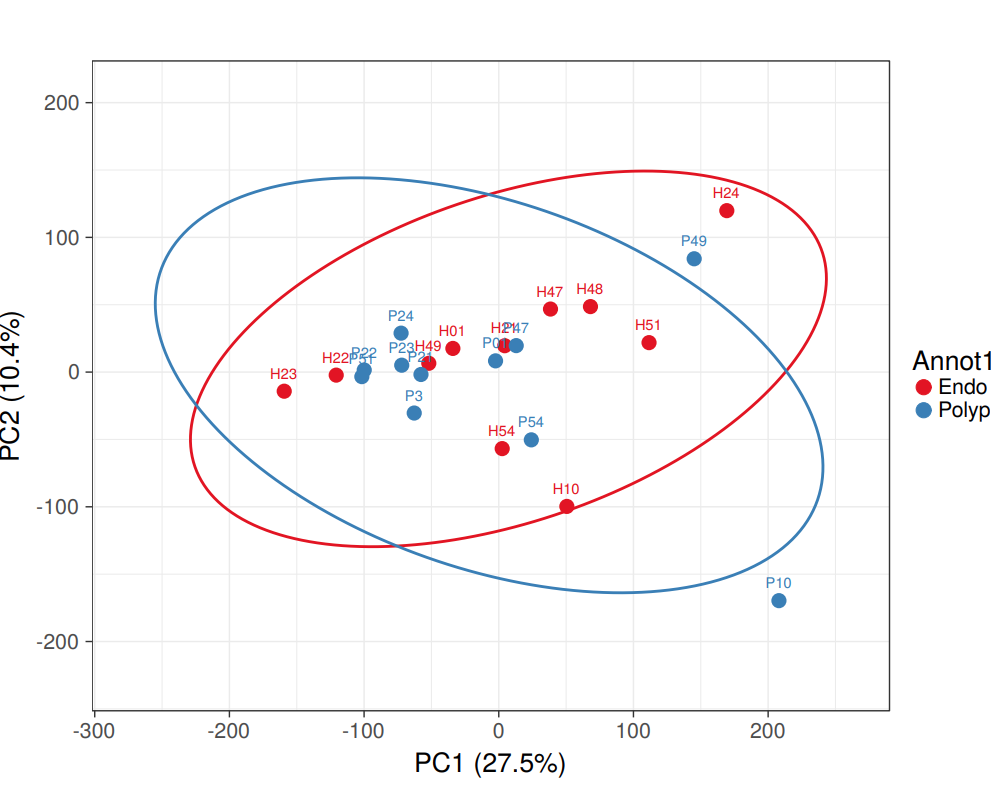

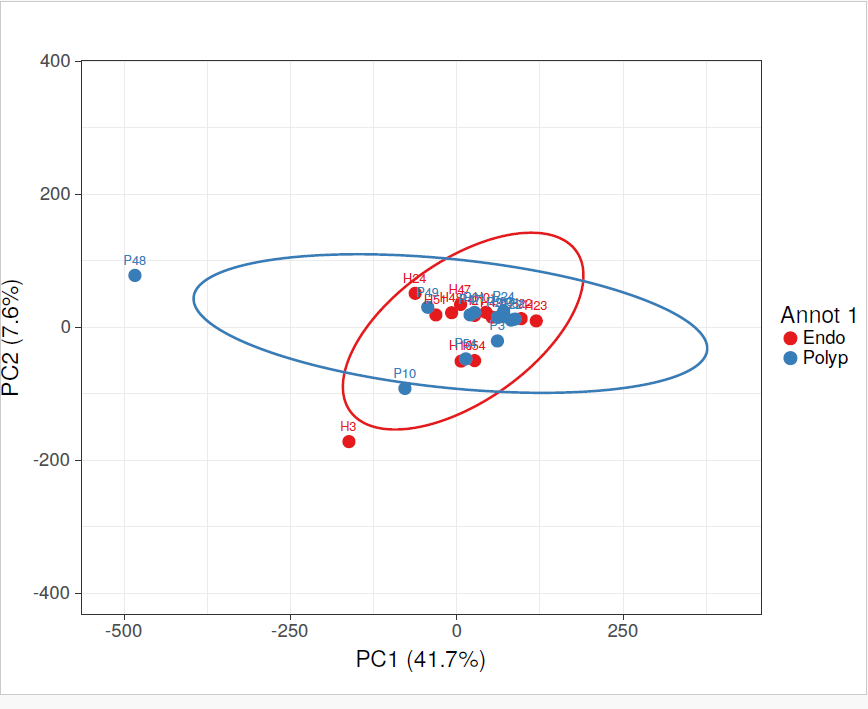
**a.	b.**

a. Principal component analysis (PCA) for differential gene expression between EPs versus adENs

b. PCA for differential gene expression between EPs versus adENs after removing outliers P48 and H3.

In the figure, EPs are marked as ‘P’ and adENs are marked as ‘H’
