## Supplementary Figure 2 for "Aberrant epithelialization: A plausible factor for the development of endometrial polyps"

**Supplementary Figure 2: Enrichment analysis of differentially expressed genes in endometrial polyps (EPs) and adjacent endometrium (adENs) in secretory versus proliferative phase**

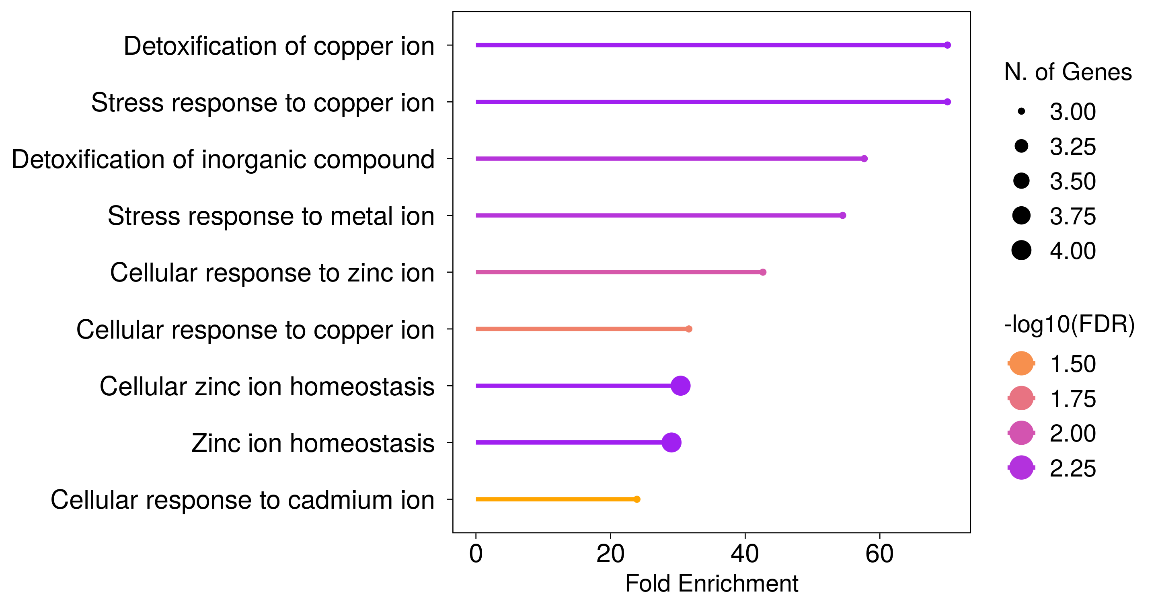
**a.	b.**

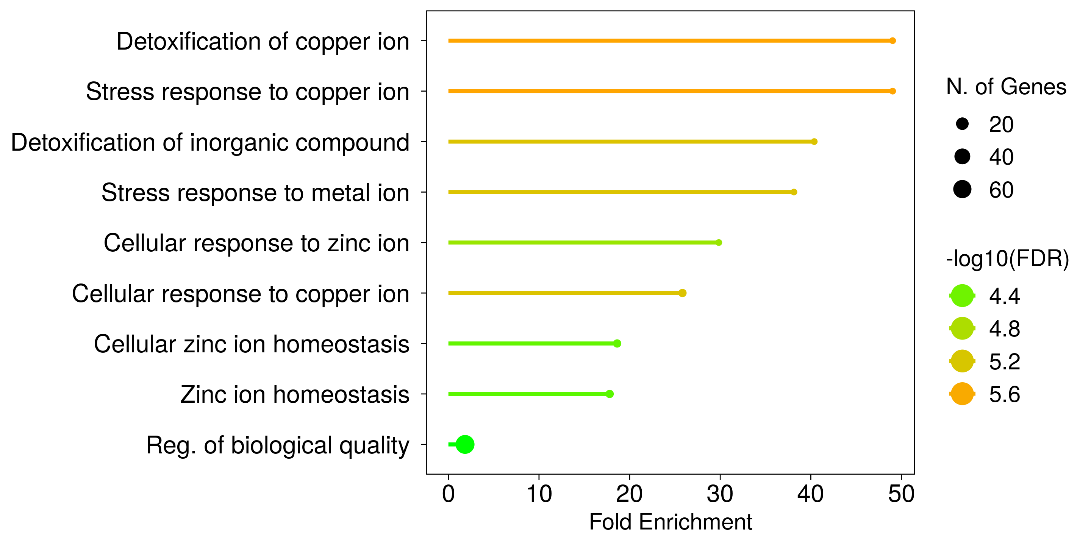

a. Biological processes involved in EPs in the secretory phases compared to the proliferative phase

b. Biological processes involved in adENs in the secretory phase compared to the proliferative phase
