## Supplementary Figure 3 for "Aberrant epithelialization: A plausible factor for the development of endometrial polyps"

**Supplementary Figure 3: Illustrative figures of histology of endometrial polyp (EP) and adjacent endometrium (adEN)**

The histology of endometrial polyps (EPs) exhibited the presence of common characteristics of EPs, such as thick-walled vascular cells, irregularly shaped glands with vacuolated epithelium and stroma with edematous-fibrotic changes.

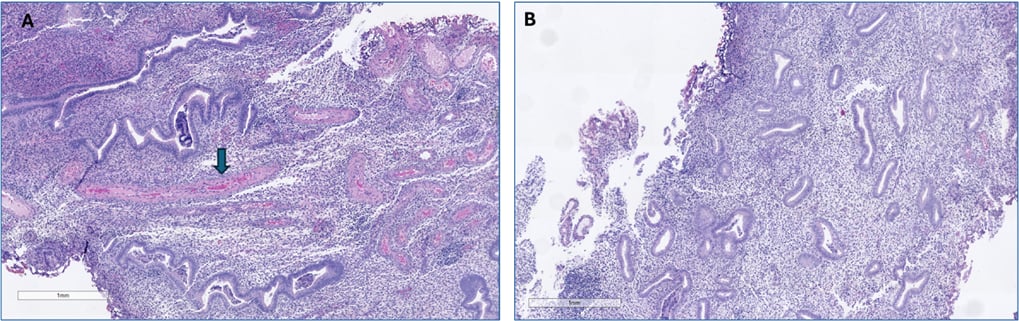

**A.** The proliferative phase EP with characteristic thick-walled blood vessels (denoted with arrow). **B.** Normal adEN tissue from the same woman. Scale bar 1 mm.
