## Supplementary Table 1 for "Aberrant epithelialization: A plausible factor for the development of endometrial polyps"

**Supplementary Table 1: Single-cell sequencing parameters and quality control parameters**

| **Samples** | **H01** | **H21** | **H23** | **H33** | **Total** | **P01** | **P16** | **P21** | **P22** | **P33** | **Total** |
| --- | --- | --- | --- | --- | --- | --- | --- | --- | --- | --- | --- |
| Estimated Number of Cells | 4092 | 1629 | 204 | 1760 | 7685 | 1961 | 2566 | 2909 | 694 | 1077 | 9207 |
| Before filtering and after a quality check | 4087 | 1556 | 199 | 1752 | 7594 | 1848 | 2431 | 2809 | 647 | 1067 | 8802 |
| After filtering (doublet removal) | 3968 | 1294 | 160 | 1589 | 7011 | 1654 | 2259 | 2605 | 565 | 992 | 8075 |
| **Groups** | **Endometrial tissues** | | | | | **Polyp tissues** | | | | | |
| **Sample** | **H01** | **H21** | **H23** | **H33** | **Average** | **P01** | **P16** | **P21** | **P22** | **P33** | **Average** |
| Median Genes per Cell | 3438 | 3869 | 1450 | 3392 | 3037.3 | 3736 | 4307 | 3693 | 2871 | 3722 | 3665.8 |
| Total Genes Detected | 29901 | 28663 | 22385 | 29109 | 27515 | 29578 | 29888 | 29307 | 25648 | 27359 | 28356 |
